# Codiversification of gut microbiota with humans

**DOI:** 10.1101/2021.10.12.462973

**Authors:** Taichi A. Suzuki, Liam Fitzstevens, Victor T. Schmidt, Hagay Enav, Kelsey Huus, Mirabeau Mbong, Bayode R. Adegbite, Jeannot F. Zinsou, Meral Esen, Thirumalaisamy P. Velavan, Ayola A. Adegnika, Le Huu Song, Timothy D. Spector, Amanda L. Muehlbauer, Nina Marchi, Ran Blekhman, Laure Ségurel, Nicholas D. Youngblut, Peter Kremsner, Ruth E. Ley

## Abstract

Some gut microbes have cospeciated with hominids, but whether they further codiversified with human populations is unclear. Here, we identify predominant gut microbial species sharing a parallel evolutionary history with human populations. Patterns of strain transfer between populations are generally consistent with an African origin, and suggest long-term vertical transmission over thousands of generations. We show the same strains also faithfully transmit between mothers and their children. Consistent with the development of intimate symbiosis, species with strongest patterns of codiversification have the smallest genomes. This study reveals long-term fidelity of gut microbiota with human populations through transmission among individuals living in close proximity. Dominance of specific strains in different populations is based in part on vertical transmission and they may provide population-specific health benefits.

**One-sentence summary:** Identification of gut microbes that codiversified with human populations.

## Main text

Coevolution, an evolutionary process involving reciprocal selection between two or more species, is more likely to occur when the partners share a parallel evolutionary history, a pattern termed codiversification (*1*). Codiversification between members of the gut microbiota and their mammalian hosts has been suggested for specific gut bacteria, based on host phylogenies matching bacterial single-gene phylogenies (*2*–*4*). Using this approach in wild apes, Moeller et al. (2016) demonstrated that several gut bacterial families codiversified with hominids over the last 15 million years (*5*). These patterns of codiversification may extend within human populations, as has been shown for one species so far. In a landmark study using multiple locus sequence typing of cultured strains, Falush et al. showed that *Helicobacter pylori*, the stomach bacterium and causative agent of gastric cancer, codiversified with its human hosts, recapitulating patterns of human migration (*6*).

The process of host-microbial codiversification often implies vertical transmission of microbes within related individuals. A recent analysis of captive apes indicates that a shared modern environment could obscure patterns of codiversification, and that geographic separation may be necessary to maintain them (*7*). Within human populations, specific gut microbes are known to transmit vertically from mothers to children (*8, 9*), which would allow for their codiversification. Methods that use multiple marker genes for inferring microbial genomes from metagenomes have revealed that a number of commensal gut microbial species also have dominant strains that differ between human populations (*10*–*13*). Although environmental factors are sufficient to drive population-level differences in strain dominance, multigenerational vertical transmission may also underlie some of these geographic patterns.

Here, we assessed codiversification between gut microbial species and their human hosts by comparing their evolutionary histories. Human- and microbial-genomes from the same individuals allowed us to assess codiversification for a suite of gut microbial species. To also gauge short-term vertical transmission, we characterized strain-sharing within mother-child pairs. After enrolling healthy adult women, with an emphasis on mothers with young children, into the study in three countries (Gabon, Vietnam, Germany), we generated chip-based genotype data for the mothers and fecal metagenomes for mothers and children (Adults/Children for Gabon: 171/144; Vietnam: 192/164; Germany: 151/78). Additionally, we generated human genotype data from Cameroon (n=115) matching to published adult fecal metagenomes (*14, 15*), and used matching human genomes and fecal metagenomes in adult subjects from the United Kingdom (n=118) (*16*). Altogether, we assessed codiversification between humans and gut microbial taxa for 747 adults and 386 children, and within generation strain transmission in 386 mother-child pairs (Fig. 1A) (Table S1&S2).

**Figure 1.**
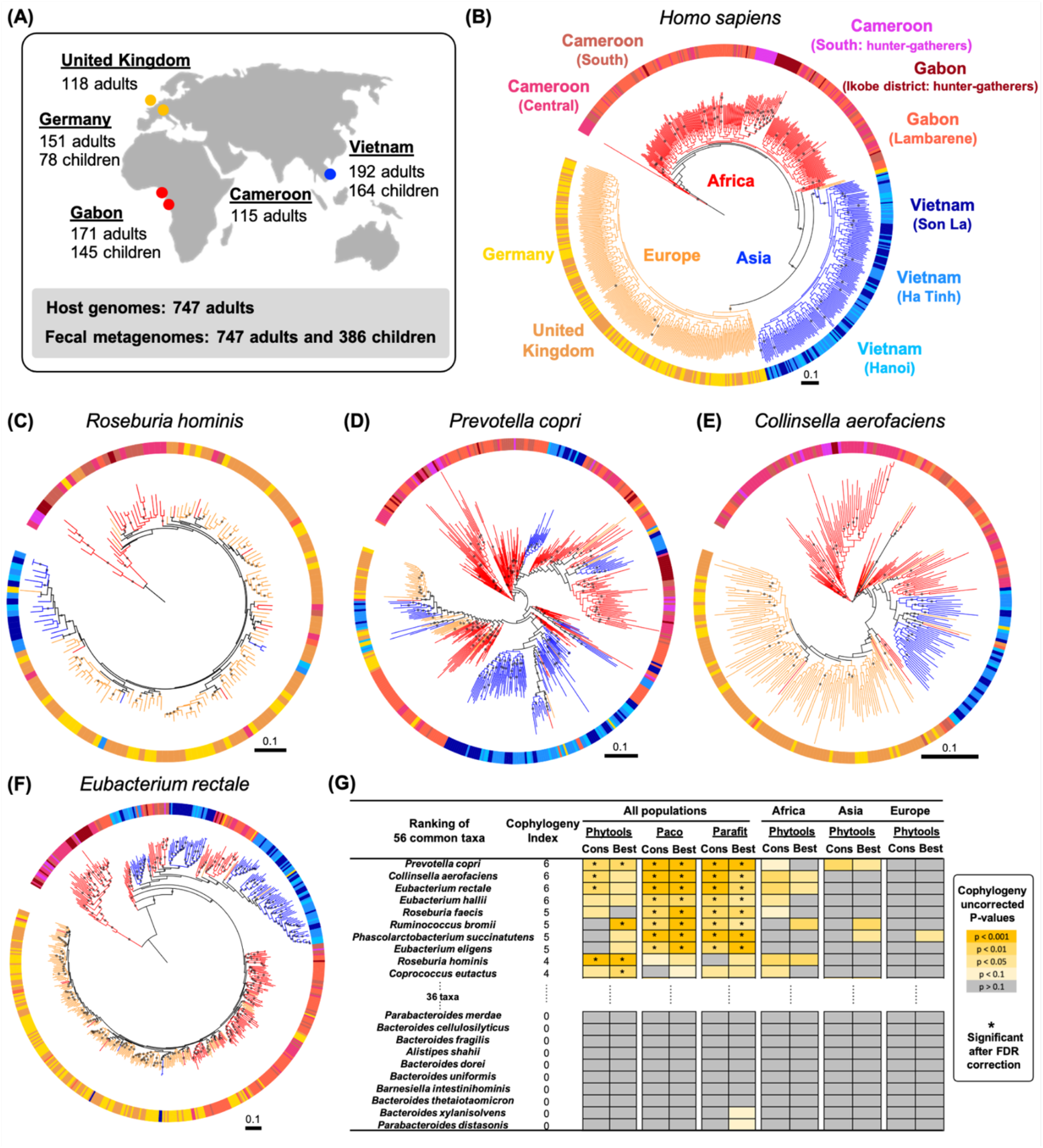
Evidence of codiversification between humans and gut microbes. (A) Study design showing sampling locations and sample sizes. (B) A maximum likelihood phylogeny of human hosts based on 32,723 SNPs. (C-F) Maximum likelihood phylogenies based on species-specific marker genes for four bacteria species that showed significant codiversification based on the strictest test (Phytools using consensus trees) after FDR correction. Bootstrap values > 50% are plotted and all phylogenies were rooted at the midpoint. The scale bars show substitutions per site. Colors of branches and outer strips correspond to sampling locations shown in panel B. (G) Codiversification test p-values based on between and within host populations for the 10 top- and bottom-ranking taxa. Cophylogeny index (CI) is based on the number of uncorrected p-values < 0.05 across six tests using all data: three tests (Phytools, Paco, and Parafit) on two types of trees (Cons: majority-rule consensus tree and Best: best maximum likelihood tree). Columns “Africa”, “Asia”, and “Europe” show within-population results in the respective group. Uncorrected p-values that remained significant after FDR correction are indicated by *. The codiversification test results for all 56 taxa with more detailed information can be found in Table S5.

A maximum likelihood phylogeny of the human hosts based on 32,723 SNPs showed human hosts clustering into three robust major groups matching their geographic origins: Africa (Gabon and Cameroon), Asia (Vietnam), and Europe (Germany and UK) (Fig. 1B). We recovered genomic information for gut microbes from metagenomes and built phylogenies using two methods: (i) species-specific marker genes using StrainPhlan3 (*10*), and (ii) metagenome-assembled-genomes (MAGs) constructed using PhyloPhlAn (*17*). Since the recovery of microbial sequences from metagenomes largely depends on a combination of read depth and the relative abundance of taxa (*10*), for the marker-based phylogenies we focused on 56 species detected in ≥100 adults with ≥10 individuals per major human group (see Methods and Table S3-S6). MAG-based trees were created for 32/56 taxa in adults. We quantified the degree of codiversification between the host and microbial trees using three different permutation-based methods: Phytools (*18*), PACo (*19*), and Parafit (*20*) (see Methods).

These analyses revealed that a subset of the gut microbiota shows significant codiversification with humans. Using our strictest criteria (Phytools using consensus trees), 12/56 species assessed show evidence of codiversification (see Methods) (Table S5). Within the 12 taxa, four remained significant for codiversification after FDR correction using the most conservative test (Fig. 1C-G). These four taxa include *Prevotella copri* (Fig. 1D) and *Eubacterium rectale* (Fig. 1F), which have previously been shown to exhibit population-specific strain diversity (*10*–*12*). For example, Tett et al. suggested that strain diversity of *P. copri* has an African origin before out-of-Africa migration events, and led to population specific strains and genes related to carbohydrate metabolism (*11*). The two others, *Roseburia hominis* (Fig. 1C) and *Collinsela aerofaciens* (Fig. 1E), are well-known carbohydrate fermenters and short-chain fatty acid producers and are linked to human health (*21, 22*).

To compare the relative degree of host-microbial codiversification among the 56 taxa using all tests, we defined a “Cophylogeny Index (CI)” as the number of uncorrected p-values < 0.05 across six tests (see Methods). Eight taxa had CIs of 5 or 6, including several *Eubacterium* spp. (Fig. 1G, Table S5). Intriguingly, *Eubacterium* spp. have been suggested to be under population-specific selection pressures due to their strong biogeographical patterns (*10, 13*). Other taxa with evidence of codiversification have been linked to host genetic variation, including *Bifidobacterium, Collinsella, Coprococcus*, and *Methanobrevibacter* (*36, 37*). Generally, gut microbes with high CIs belonged to the Firmicutes phylum (Fig 2A). In accord, strains within species of Firmicutes are reported as more geographically restricted compared to those of other phyla (*7*, but see *12*). In contrast, taxa with low CIs belonged to the Bacteroidetes phylum, including several species of *Bacteroides* and *Parabacteroides* (Fig. 1G, Fig. 2A Table S5), which are known to survive in environments such as water (*23*). Applying the same tests to trees constructed with MAGs resulted in consistent results (Table S7). Thus, our results identify a subset of gut microbes whose strain phylogenies are strongly indicative of codiversification with their human hosts.

**Figure 2.**
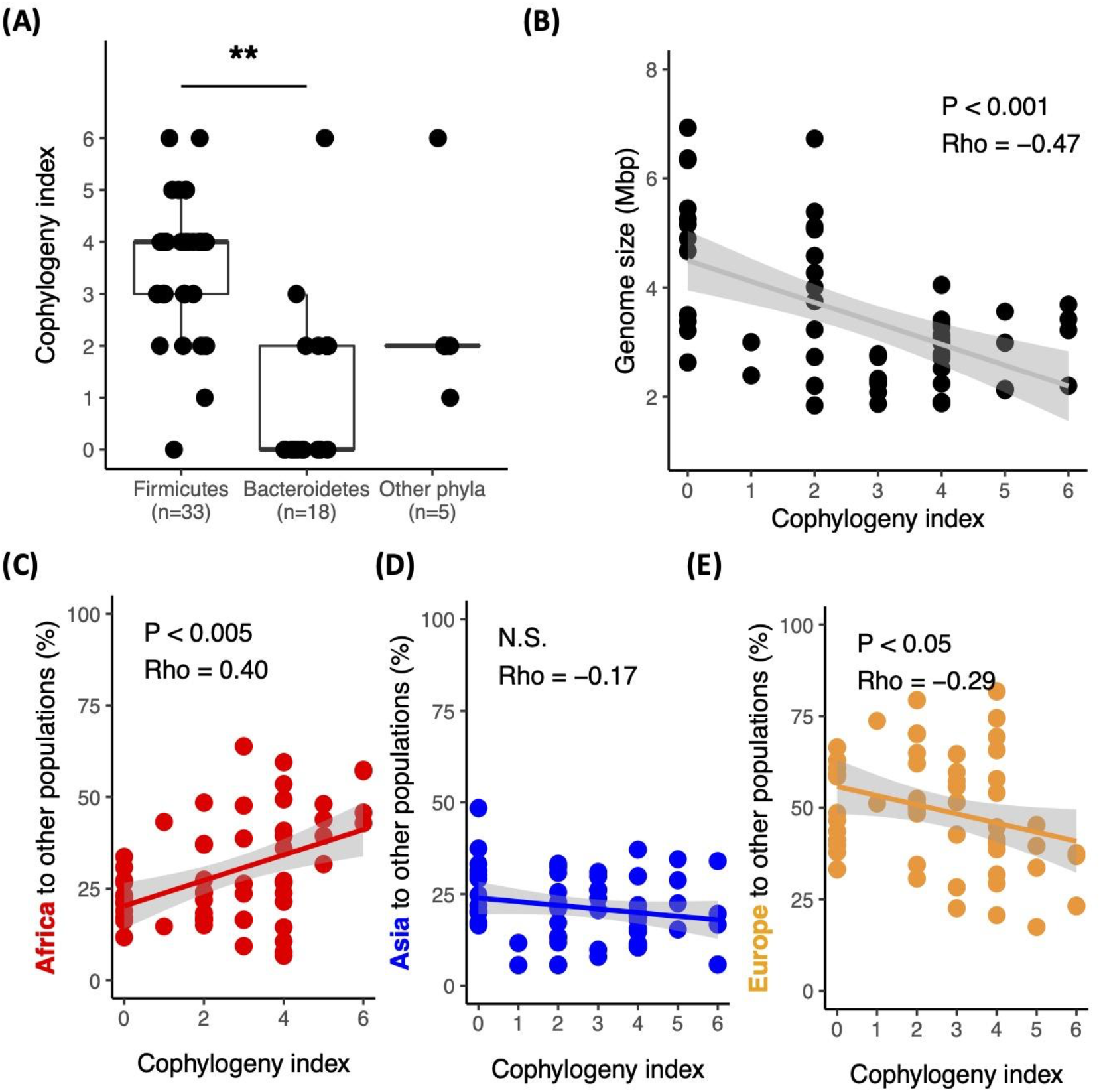
Taxonomy, genome size, and transfer events are related to host-microbial codiversification. (A) CI values by phylum. The P-value is based on the Wilcoxon rank sum test. (B) Correlation between genome size and CI. Correlations between CI and the proportions of transfer events from (C) Africa, (D) Asia, and (E) Europe.The results are based on stochastic character mapping. Each dot represents a microbial species. All correlations are based on Spearman’s rho. The 95% confidence intervals are shaded in gray. ** = p < 5×10^-5^ ; N.S., non significant (p > 0.10).

We observed a strong negative correlation between CI and genome size (Fig. 2B). Seven out of the eight genera had a negative relationship between CI and genome size independently (Binomial test p = 0.035, Fig. S1). Strict vertical transmission of symbionts in other hosts, such as insects, is associated with genome reduction of the microbe resulting from reduced purifying selection (*24*). Our results provide the first evidence in mammals that a similar process of host adaptation may be occurring through genome reduction.

It is well established that modern humans originated in Africa, and therefore long-term vertical transmission of microbial taxa would be expected to result in out-of-Africa patterns. To test for this, we quantified the number and direction of host-region-switch events by applying stochastic character mapping for all 56 microbial phylogenies (marker trees, Fig. 2C-E, Fig. S2A). We found that taxa with high CIs exhibited significantly more transfer events from Africa to other regions, compared to taxa with low CIs (rho = 0.40, p = 0.003) (Fig. 2C). Transfer events from Asia to other regions were the same for the top and bottom CI taxa (rho = -0.17, p = 0.22) (Fig. 2D). In contrast, taxa with low CIs had higher proportions of transfer events from Europe to other regions, compared to high-CI taxa (rho = -0.29, p = 0.029) (Fig. 2E). MAG-based trees resulted in similar patterns (Fig. S2B). Overall, the results are consistent with an African origin for high CI taxa.

To corroborate the out-of-Africa pattern with independent data, we used 1,219 public human gut metagenomes (Fig. 3A, Table S8&9). Strains derived from the public metagenomes generally clustered with those derived from our samples based on the sampling locations (Fig. 3B-F, Fig. S3), and the resulting phylogenies were consistent with those of previous studies (*10, 11, 13*). We observed the same out-of-Africa strain transfer patterns for the top CI-taxa using the public metagenomes with (Fig. 1&3, Fig. S2C) or without the five populations used in the initial analyses (Fig. S2D). We also included primate fecal metagenome data (n=107), and for high CI-species, strains from primate hosts tended to be the most early diverging strains in relation to all human strains (Fig. 3, Fig. S3). In contrast, low-CI taxa had primate strains nested within human strains (Fig. S4). Overall, the results are consistent with human gut microbial taxa originating from ancestors common to hominids (*5*), and extend this process into human populations as modern humans migrated from Africa.

**Figure 3.**
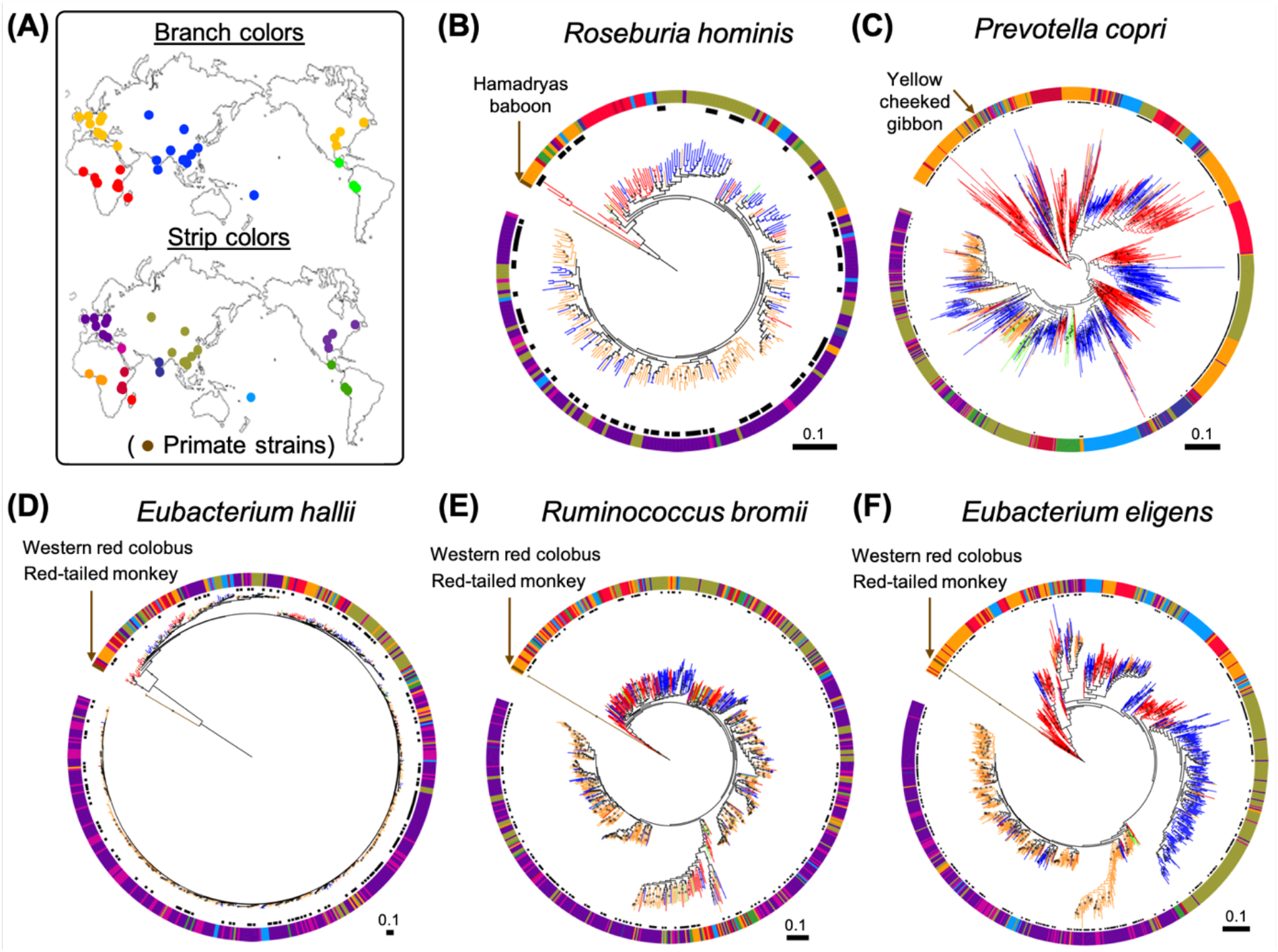
Publicly available fecal metagenomes of humans and primates support the original findings. (A) Map showing the source of fecal metagenomes used in B-F. The colors of the sampling location dots on the map correspond to colors used in B-F for the branches and outer strips. Colors are human genetic structures estimated from sampling locations based on data in Duda and Zrzavy et al. (*27*) (Table S8&9). (B-F) Maximum likelihood phylogenies of five out of the 10 top-ranking taxa constructed using data generated here (black dots next to the outer strip) combined with data from human and primate public metagenomes. All primate strains in the trees are labeled. Bootstrap values > 50% are plotted. All phylogenies were rooted at the midpoint. The scales are substitutions per site.

Our inference of strain transfer events is based on patterns within the phylogenies for present-day strains, yet ancient DNA analysis can provide a snapshot directly from the past. Using high-quality ancient MAGs derived for five gut microbial species recovered from 1,000 - 2,000 year old paleofeces of Native North American tribes (*25*), we observed that four out of the five taxa (the exception is *Escherichia coli*) were most closely related to strains from modern East Asians (Fig. S5), consistent with the known migration history of the Americas (*26*).

Next, we tested for evidence of codiversification in the child microbiomes using the mothers’ genotypes. *P. copri* showed evidence of codiversification (p < 0.05) in both adults (Fig. 1D, Table S5) and children (Fig. S6, Table S6). In addition to *P. copri*, we observed *Bifidobacterium* species with high CIs in children. Among the 20 common taxa tested (see Methods), 8/20 showed significance (p < 0.05), including all four *Bifidobacterium* species (based on the strictest test), and three remained significant after FDR correction (*B. bifidum, B. longum*, and *B. breve*; Fig. S6). Using MAG-based trees, we tested 11 taxa: *B. longum* also showed significance based on the strictest test (Table S10). As with adults, child taxa with highest CIs tended to have transfer events from Africa to the rest of the populations (p = 0.052 Fig. S2E). Our results extend those of previous studies, where the bacterial family Bifidobacteriaceae exhibited codiversification with hominid species (*5*).

Signals of codiversification extended within populations for both adults and children. Within-population tests had smaller sample sizes and the signal of codiversification tended to be weaker: none of the p-values remained significant after FDR correction for adults (Fig. 1G, Table S11), and only one taxon remained significant in children (*Bifidobacterium bifidum* within Africa, Table S6). The patterns are nevertheless consistent: many of the same taxa with high-CI also showed evidence of codiversification within populations, particularly for Africa (Fig. 1G). For example, *P. copri* showed evidence of codiversification within-populations in adults (Africa: p = 0.089 and Asia: p = 0.0027; for children p = 0.094). Host population differences in diet likely affect the abundance and diversity of *P. copri* strains (*11*). Our results between- and within-populations support long-term vertical transmission contributing to the observed pattern as well.

Taxa that are transmitted vertically over thousands of generations are expected to be shared in mother-child pairs today. We used two independent approaches to search for shared strains in mother-child pairs: inStrain (*28*), a tool based on population-level nucleotide strain diversity (popANI), and SynTracker (*29*), based on the synteny of long-stretches of DNA. We identified bacterial species with strains more similar between mothers and their own children compared to unrelated children. Of the 63 taxa (adult and child taxa combined) assessed for codiversification, 20 were tested for strain-sharing events between related and unrelated mother-child pairs. Of these, notably, strains of *P. copri* and other *Prevotella* spp. were shared between mothers and their children using two independent methods (Fig. 4A, Fig S7&S8). *E. rectale* also showed evidence of mother-child strain sharing (Fig. 4A&S7). These results are consistent with the observed codiversification patterns (Fig. 1, Fig. 3C, Fig. S6).

**Figure 4.**
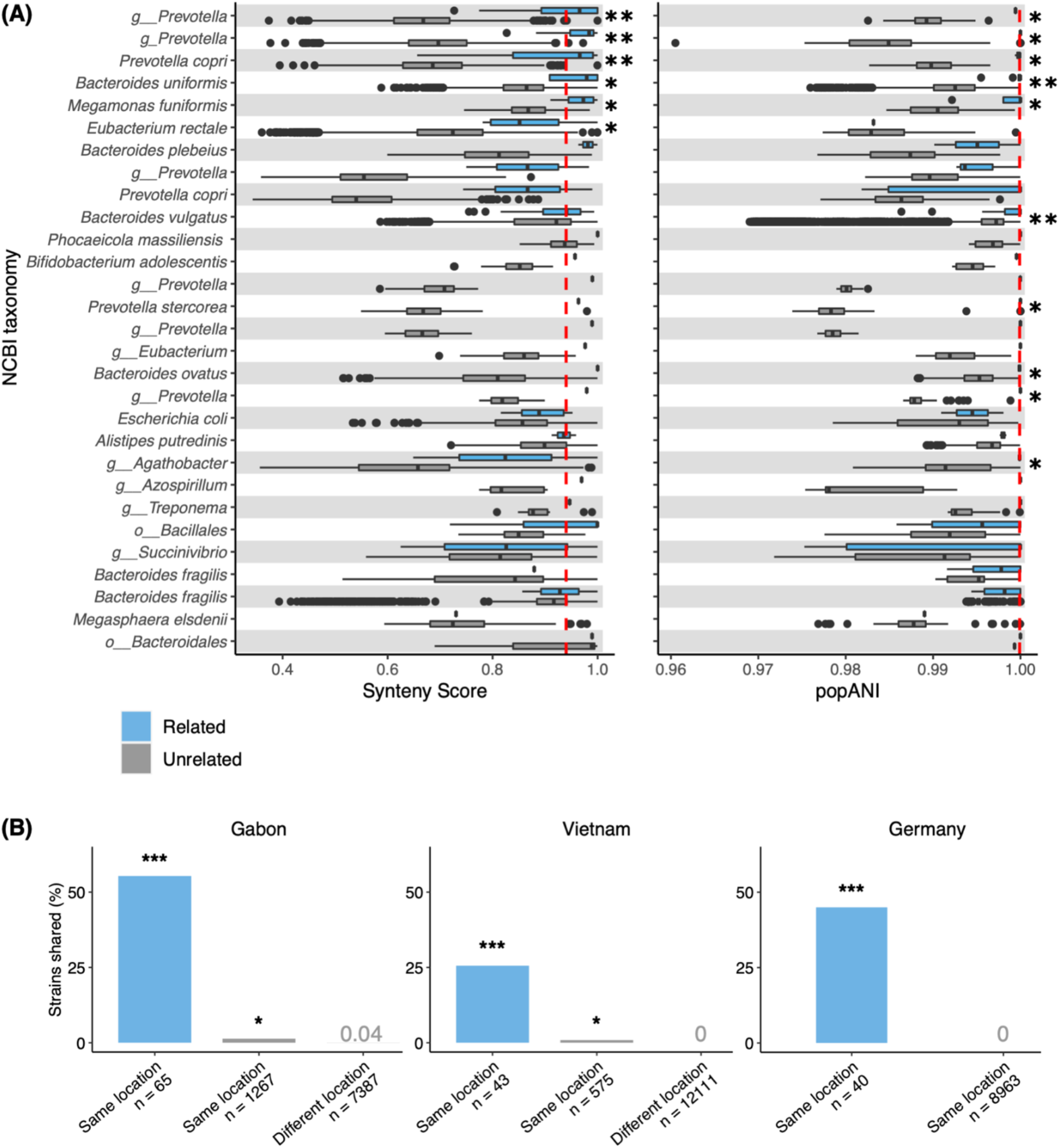
Strain sharing assessed within related and unrelated mother-child pairs in Gabon, Vietnam, and Germany. (A) The relatedness of strains per species within related and unrelated mother-child pairs, using SynTracker (left) and inStrain (right). Species listed are those identified in both related and unrelated mother-child pairs. Dashed red lines indicate the pairwise comparison thresholds for strain sharing events (0.96 for synteny; 0.99999 for popANI). The full list of taxa tested in each method is shown in Fig. S7&8. (B) Strain sharing within related and unrelated mother-child pairs by country using inStrain. The percentage of total strain comparisons (“n”) identified as strain sharing events between related mother-child pairs, unrelated mother-child pairs sampled in the same location, and unrelated mother-child pairs sampled in different locations. Stars correspond to q-values (corrected Wilcoxon-Mann-Whitney test for Fig. 4A and corrected hypergeometric test for Fig. 4B; * = p < 0.05; ** = p < 5×10^-5^; *** = p < 5×10^-25^; Table S12). Comparisons were calculated with samples collected from 6 locations in Gabon (average inter-location distance = 114km), 4 locations in Vietnam (average inter-location distance = 238km) and 1 location in Germany (Table S13). The phyla comprising the strain sharing events shown here are described in Fig. S9.

In addition to mother-child strain transmission, we also observed strain sharing between mothers and children who were not their own, particularly for individuals sampled in the same location (Fig. 4B). Higher rates of transmission among socially engaging individuals (that are often genetically related) are known in group-living species such as bees (*30*), mice (*31*), and great apes (*32, 33*), and this can provide signals of codiversification at the population-level but weak signals of vertical transmission between mother-child pairs. Similarly, for taxa where we find no evidence of codiversification but evidence of vertical transmission, such as *Bacteroides uniformis* (Fig. 1G, Fig. 4B), this pattern may be due to changes in lifestyle across time (*e*.*g*., greater sanitation or isolation reducing group-level transmission).

This work yielded a first list of taxa that codiversified with humans, yet more likely remain to be identified. Indeed, less prevalent and/or low-abundance taxa will have been missed due to low coverage. The taxa most impacted may be those at low relative abundance in any one of the major human groups (*i*.*e*., Africa, Asia, and Europe): such taxa would be excluded in this study. In addition, even though we could detect several strains per species per subject using SynTracker and inStrain, we only tested for codiversification for the more dominant strains because marker-based approaches extract the dominant strain per microbial species per individual (*10*), and the MAG-based approach could not distinguish closely related genomes. Targeted methods, such as genome enrichment or isolation of specific microbial taxa, could expand the list of taxa showing patterns of codiversification.

In conclusion, we have identified common members of the human gut microbiota that show evidence of codiversification with humans. Our results suggest that these species have transferred within related individuals living in proximity for thousands of generations. These findings extend previous observations of cospeciated bacteria with humans from our hominid ancestors: they further codiversified within human populations. Gut microbiota that codiversified with early modern humans as they migrated to populate the world were exposed to new environments and may have participated in the processes of local adaptation in humans (*35*). Because some of the codiversified taxa have been linked to a variety of diseases in humans (*6, 39*), functional differences among codiversified strains may contribute to discrepancies between studies addressing the role of microbiome in physiology in different populations (*12*). The list of gut microbial taxa provided here should motivate future studies to investigate the roles of codiversified strains in human health and host-microbial coevolution.

## Acknowledgements

We thank Nguyen Thu Ha, Anne Pfleiderer, Emily Cosgrove, Andrew Clark, Aleksander Kostic, Marla Taylor, Native American tribes (Mike Bremer, Joseph Aguilar, Juana Charlie, Reylynne Williams, Barnaby Lewis, Shane Anton, Angela Garcia-Lewis, and Bruce Bernstein), and members of the Department of Microbiome Science.

## Data and material availability

The raw sequence data and MAGs are available from the European Nucleotide Archive under the study accession number PRJEB46788. All sample metadata used in this study are provided in Supplementary information. The code used in data analysis is outlined on GitHub at https://github.com/leylabmpi/codiversification. The raw metagenome sequence data are available from the European Nucleotide Archive under the study accession numbers PRJEB47532, PRJEB47543, PRJEB40256, PRJEB9584, PRJEB32731, PRJEB27005, PRJEB30834, and PRJEB46788.

