## Supplemental_Methods_Figures_S1_S9 for "Codiversification of gut microbiota with humans"

### Material and Methods

Recruitment of Participants

Five hundred and fourteen adult women were recruited for the study from three countries: Gabon (n = 171, average age = 27 years old), Vietnam (n = 192, average age = 28 years old), and Germany (n = 151, average age = 29 years old) (see Table S1 for detail). Inclusion criteria included: women between the ages of 18 and 40; no severe allergy to dairy products; not currently self-reported as pregnant; no known underlying medical conditions; local residency. Three hundred and eighty six children of included participant mothers were recruited: Gabon (n = 144, average age = 8.0 months), Vietnam (n = 164, average age = 6.7 months), and Germany (n = 78, average age = 7.7 months) (see Table S2 for detail). Participants were screened for the above criteria by field staff and/or physicians upon arrival at the participating clinics, and all protocols, risks, and study motivations were carefully explained to participants in local languages. All participants provided informed consent for participation in the study for themselves, and as applicable, for their child. All human research was approved by local ethical committees. The study protocols for this study are as follows: Gabon, issuing authority: Comité National d'Ethique, Protocol number: N0025/2017/PR/SG/CNE; Vietnam, issuing authority: Scientific Ethics Review Committee and 108 Military Central Hospital, Protocol number: 108MCH/RES/VGCARE-03-16012018; Germany, issuing authority: Ethik-Kommission an der Medizinischen Fakultät der Eberhard-Karls-Universität und am Universitätsklinikum Tübingen. 529/2018BO1.

Generation of gut metagenome data

Participants used Fe-Col collection kits (Alpha Labs, Eastleigh, UK) to collect stool from themselves and/or their child. Samples were frozen on dry ice within 8 hours of collection and transported on dry ice for storage at -80°C. Genomic DNA was extracted from frozen stool using PowerSoil DNA extraction kits (Qiagen, Hilden, Germany). Metagenomic library preparation was performed as per Youngblut et al. (2020)(*1*) with slight modifications. Briefly, 1ng of purified gDNA was used in a Nextera (Illumina, San Diago, USA) Tn5 tagmentation reaction to fragment and ligate adaptors in a single reaction, followed by a 14 cycle PCR to add sample specific barcodes. Libraries were purified using Mag-Bind TotalPure NGS beads (Omega Biotech, Norcross, USA), pooled, and quantified. Size selection (400-700bp) was performed on a BluePippin (Sage Science, Beverly, USA). Libraries were concentrated and further purified as needed using DNA Clean & Concentrator-5 (Zymo Research, Irvine, USA). Sequencing was conducted on a HiSeq 3000 System (Illumina, San Diago, USA) with 150 paired-end sequencing. We used a quality control pipeline described in Youngblut et al. (2020)(*1*). Briefly, adapter trimming and quality control filtering was conducted using Skewer0.2.2 and bbtools "bbduk" command. Reads mapping to the human genome (GRCh37/hg19) were filtered using bbtools "bbmap" command. The read quality was assessed using Fastqc 0.11.7 and multiQC 1.5a. The same pipeline was applied to all publicly available metagenomes that were used in this study including the previously published metagenomes from Cameroon (PRJEB27005 and PRJEB30834) (*2*, *3*) and the UK (PRJEB13747) (*4*) and the 1326 public metagenomes from various studies (see below).

<u>Metagenome-assembled genome (MAG) generation and analysis</u>

Paired-end reads were subsampled to ≤20 million per sample with seqtk 1.3. We used metaSPAdes 3.12.0 for per-metagenome *de novo* assemblies. Contig binning with differential coverage was conducted with three binners: MetaBAT 2.15, MaxBin 2.2.7, and VAMB 3.0.2. We ran each binner 3 times with different parameter sets (MetaBAT: [-prob_threshold 0.6, -

prob_threshold 0.7, -prob_threshold 0.8], MaxBin: [--maxP 92 --maxEdges 150, --maxP 94 --maxEdges 325, --maxP 97 --maxEdges 500], VAMB: [-l 24 -n 384 384, -l 32 -n 512 512, -l 40 -n 768 768]) for a total of 9 binning methods applied to each metagenome assembly. Only contigs ≥1.5 kbp were used for binning. Per-metagenome differential coverage binning with coverage calculated from all other genomes would require an excessive number of read mapping jobs (*i.e.,* all pairwise combinations of metagenomes), so we utilized a subsampling approach. Specifically, for each focal metagenome, we randomly selected 40 metagenomes, with the focal metagenome always included in the selection. Reads from the focal metagenome were only mapped to this metagenome subset in order to assess differential coverage for the focal metagenome. Bowtie 2.4.1 was used for mapping reads to contigs. To reduce biases in coverage estimated resulting from varying sampling depths among metagenomes, we subsampled to ≤5 million paired-end reads prior to mapping. We used DAS-Tool (1.1.2 for the current study and Lokmer et al. 2019 dataset; 1.1.3 for the Even et al. 2021 dataset) to select the highest quality, non-redundant contig bins (MAGs) from all 9 binning methods, with quality based on CheckM 1.1.3-estimated completeness and contamination.

The raw MAGs were filtered by CheckM-estimated completeness (≥ 50%) and contamination (< 5%) and then dereplicated at 99.9 and 95% ANI with dRep (3.2.0 for current study; 3.2.2. for Lokmer et al. 2019 and Even et al. 2021). Prodigal 2.6.3 was used for gene prediction. Multi-locus genome phylogenies were inferred via PhyloPhlAn 3.0.2, with DIAMOND 2.0.9 used for phylogenetic marker homology searches, MAFFT 7.480 used for the multiple sequence alignment, FastTree 2.1.10 used for initial tree inference, and RAxML 8.2.12 used for final tree inference (starting from the phylogeny inferred from FastTree). Outgroups for each clade-level phylogeny were automatically selected from sister clades (*e.g.,* sister genus). Specifically, the GTDB Release 95 genome metadata was used to identify genomes in sister clades, and the outgroups were selected with preference given to genomes with the highest CheckM-estimated quality (*i.e.,* completeness and contamination).

<u>Mother-child strain sharing analysis</u>

We used CheckM and dRep to dereplicate 6,234 filtered MAGs assembled from mother-child metagenomes at 95% ANI and to identify 796 species representative genomes (SRGs). Species were assigned to both GTDB and NCBI using gtdb_to_taxdump v0.1.6 (*5*). We mapped metagenome reads from 386 mother-child pairs to the SRGs with inStrain 1.5.3's "profile" function (--min_cov 5 --min_freq 0.05 --min_genome_coverage 0.1 --min_read_ani 0.95 --database_mode --skip_plot_generation) and used the "compare" function to calculate popANI and percent_genome_compared (--min_cov 5 --min_freq 0.05 --ani_threshold 0.99999 --database_mode --skip_plot_generation). Pairwise comparisons with popANI ≥99.999% and percent_genome_compared ≥50% were identified as strain sharing events and percent_genome_compared ≥50% was used. To test whether there are significant strain-sharing events within related mother-child pairs compared to within unrelated mother-child pairs more than expected by chance, we used the hypergemetric distribution R function "phyper", with correction for multiple comparisons using the R function "p.adjust" (*--method* = "BH"; Table S12).

We used SynTracker (*6*) to quantify the relatedness of strains in related and unrelated mother-child pairs using the 6,234 filtered MAGs, with each SRG as a reference genome, with default parameters (--perc_identity = 97, --qcov_hsp_perc=70, --maxSep=15, --maxGap = 15). Average pairwise synteny scores were calculated by randomly selecting 30 homologous 5kbp regions/pairwise. Significance tests for the synteny scores and popANI of strains between related and unrelated mother-child pairs were performed using the R function "wilcox.test" (--

*alternative* = less). Multiple testing correction was performed using the "p.adjust" function (--*method* = "BH"). Note that unlike the marker-based phylogenies described below (*i.e.,* studying the dominant strain per individual), results from inStrain and SynTracker represent multiple strains per individual.

Data generation of human genotype data

We generated human genotype data from the participants in four countries (Gabon, Vietnam, and Germany, and Cameroon) and used published genotype data from the United Kingdom (*4*). In Gabon, Vietnam, and Germany, we collected saliva samples from each participant using Saliva DNA Collection and Preservation Devices (Norgen, Thorold, Canada). Genomic DNA from saliva was extracted using PowerSoil DNA extraction kits (Qiagen, Hilden, Germany). DNA was analyzed using the Infinium Global Screening Array (Illumina, SanDiego, USA) and genotyped at the University Hospital of Bonn, Life & Brain Research Centre. For Cameroon participants, host genotype data was generated for individuals that were collected in 2013 and 2017 with published gut metagenomes (*2*, *3*). For each participant, approximately 2 ml of saliva was collected in a 15 ml falcon tube, to which we added the same volume of a homemade buffer (5mM TRIS, 5mM EDTA, 5mM sucrose, 10mM NaCl and 1% SDS, pH=8). DNA was then extracted following the procedure from (*7*). For samples collected in 2013, the extracted DNA was then genotyped on an Illumina Omni2.5 genotyping array at the University of Chicago Genomics Core, in Chicago, USA. For samples collected in 2017, after quantification using a PicoGreen assay and qPCR assays to check for amplification and clustering, extracted DNA was genotyped on an Illumina Multi-Ethnic Global array at the University of Minnesota Genomics Center.

Using the GenomeStudio software, we excluded individuals with a call rate below 0.95, as well as markers that failed genotyping, present on non-autosomal chromosomes, with a call rate below 95% or a cluster separation below 0.3. For the previously published genotype data from the UK (*4*), we excluded close relatives (*i.e.*, one individual per twin) and individuals that do not have paired fecal metagenomes. Finally, genome assembly versions were converted to GRCh38 using UCSC Genome Browser LiftOver and all datasets were merged in PLINK v1.9. After merging, further quality control was conducted where variants that are non-biallelic, minor allele frequency of $< 0.05$, missing call rates of $> 0.9$, and deviated from Hardy-Weinberg equilibrium ($P < 10^{-5}$) were removed. This resulted in 32,723 SNPs that overlapped across all five datasets.

<u>Human phylogenetic inference from SNP data</u>

We created 100 maximum likelihood trees in SNPhylo (version 20180901) (*8*) including 747 individuals using 32,723 SNPs, which were further filtered to remove uninformative SNPs based on the following parameters (ld_threshold = 0.1, maf_threshold = 0.05, and missing_rate = 0.1). This resulted in around 10,000 SNPs to create the host phylogeny (each bootstrapped tree uses a slightly different number of SNPs). We picked the best tree out of the 100 trees based on the maximum likelihood score and plotted the bootstrap values (referred as "HostBestTree") using "plotBS" in phangorn R package. To account for branches that have low bootstrap support, a majority-rule consensus tree was also created where the best tree with branches with bootstrap values < 50% were collapsed (referred as "HostConsTree") using the function "as.polytomy" in ggtree R package. The host tree was rooted by midpoint. Final trees were annotated in ITOL v6.

<u>Microbial phylogenetic inference from StrainPhlAn data</u>

We created microbial phylogenies from adult and child metagenomes using StrainPhlAn v3.0 (*9*). First, we picked the top 100 prevalent taxa in metagenomic samples of adults (n=747) and children (n=386) using MetaPhlAn3 (v3.0.1) using the following parameters (--tax_lev a --min_cu_len 2000). The taxonomy is based on NCBI. Next, we used StrainPhlan3 to generate microbial phylogenies using the following parameters: samples2markers.py (--breadth_threshold 80) and strainphlan.py (--phylophlan_mode accurate --marker_in_n_samples 10 --sample_with_n_markers 10). To allow for between-population comparisons, we selected taxa in adults that represent a total of ≥ 100 individuals and ≥ 10 individuals each from major human grouping; Africa (Cameroon and Gabon), Asia (Vietnam), and Europe (Germany and UK) which resulted in 56 taxa (Table S3&S5). For child samples, we picked the top 20 taxa that had the largest trees due to the smaller sample size (Table S4&S6). Similar to host trees, the best tree out of 100 trees based on the maximum likelihood score was selected (referred as "BacBestTree"), and a majority rule consensus tree was created (referred to as "BacConsTree"). We chose StrainPhlan3 for the main method to create strain-level phylogenies because it is suited for creating phylogenetic trees of the most dominant strain per individual. All trees generated by StrainPhlAn3 are rooted by midpoint. All MAG-based trees are rooted by an outgroup. Final trees were annotated in ITOL v6.

Public metagenomes and phylogenetic trees

We downloaded and curated accessible public metagenomes at the time of the study (Table S8&9). We retained stool samples with no disease status (healthy or control group), no current use of antibiotics, and age ranging from 12-65 years old (age mean ± s.d. = 36.7 ± 13.6 years old). All samples that indicated non-local individuals (*e.g.,* travelers) were also excluded.

For visualization purposes, the collection locality of the individual was used as a proxy for host genetic groups based on Duda and Zrzavy (2016) (*10*). The assignment of genetic structure and color used in Fig. 2 can be found in Table S8. When samples exceeded 100 individuals per population per study, we randomly selected 100 individuals to reduce the sampling bias. Samples were quality filtered using the same QC pipeline described above. When multiple samples were available per individual, we combined the reads. Using StrainPhlAn3 with the same parameters described above, we were able to extract marker genes from fecal metagenomes derived from 1219 human individuals representing 19 countries in 21 studies(*4*, *11–30*), and 107 primate individuals representing 27 species of primates from 12 countries in 2 studies (*31*, *32*). The number of sequence reads and detailed sample information for all public metagenomes used in this study are shown in Table S9. Phylogenies for the 10 top-ranking and bottom-ranking taxa were created using StrainPhlAn3 with the same parameters described above. High-quality ancient MAGs from paleofeces described in (*33*) were shared by Alex Kostic with support from the Peabody Museum. Six ancient MAGs that were identified at the species level were added to the microbial phylogenies as reference genomes in StrainPhlAn3 (Fig. S5). All trees were annotated in ITOL v6.

<u>Cophylogeny statistics</u>

We used three different methods to quantify the degree of codiversification between the host tree and the microbial tree: (i) function “cospeciation” in phytools R package, which is based on Robinson-Foulds distance (referred as “Phytools”)(*34*); (ii) function “parafit” in the ape R package, which is based on principle coordinate analyses and euclidean distance (referred as “Parafit”)(*35*); and the function “PACo” in paco R package, which is based on principle coordinate analysis and procrustes distance (referred as “PACo”)(*36*). A recent study noted that

PACo and Parafit tests were not designed for or tested with a large number of host-microbiome relationships (*37*). Thus, for PACo and Parafit, we collapsed highly similar microbial strains as the same strain using the function "cutreeDynamic" in dynamicTreeCut R package with the following parameters: cutHeight = 0.99 and minClusterSize = 2. This was necessary to produce meaningful p-values because all taxa resulted in p-values < 0.001 when one-to-one pairs of host-microbe pairs were considered without collapsing the similar strains. We only used the results of PACo and Parafit to calculate the Cophylogeny Index (CI, see below), and we mainly report results derived from the strictest test, Phytools, in the main text. All tests were conducted with 999 permutations. We repeated each test 10 times (each test with 999 permutations) and calculated the mean and standard deviations of p-values. To account for the uncertainty of phylogenetic topologies, we applied the three codiversification tests on majority rule consensus trees, where branches with bootstrap values less than 50% were collapsed (HostConsTree vs BacConsTree), and on best trees based on the maximum likelihood score (HostBestTree vs BacBestTree). Overall, the number of strains represented in a microbial tree did not bias the three codiversification test results (Table S14). Furthermore, p-values from the three codiversification tests tended to yield similar correlations to each other based on Spearman's rho, regardless of the microbial tree type (Table S15). The mean and standard deviations of test statistics are reported in Table S16.

#### Cophylogeny Index

We summarized the overall support of codiversification by calculating CI, the number of uncorrected p-value < 0.05 across six tests (*i.e.*, three codiversification tests on two tree types). The motivation to use uncorrected p-values for the index over FDR-corrected p-values is that the index will not change due to the number of taxa tested. The CI is useful to quantify the relative difference in the degree of cophylogeny summarizing the results from multiple tests. The probability of 12 out of 56 taxa having a significant p-value for the strictest test (Phytools with consensus trees) is 0.0054. The calculation was based on binomial probability (equation below), where X = 12, n = 56, and p = 0.1 (false positive rate, one in ten tests are significant by chance):

$$P(X) = \frac{n!}{X!(n-X)!} \cdot p^X \cdot (1-p)^{n-X}$$

Similarly, we also calculated the probability of a given taxon to show significance across all three tests. Although six tests go into the CI, we calculated the probability of three tests using consensus trees to be conservative because two tree types (HostConsTree vs BacConsTree and HostBestTree vs BacBestTree) within tests are highly correlated. If we conservatively assume that at least 6 out of 56 taxa show significance due to chance in a given test (probability of 0.060 using the equation above), the probability of the same taxa showing significance across three different tests is 0.0012, where X = 3, n = 3, and p = 6/56, or P = $(6/56)^3$. The full results of all 56 adult taxa and 20 child taxa can be found in Table S5 and Table S6.

<u>Cophylogeny: children</u>

We tested host-microbial codiversification in both adults and children. In the case of children, we used the host tree based on the mother's genotype and microbial tree based on the child's metagenome and tested for codiversification using Phytools. Similarly, we tested codiversification within the three continents independently using Phytools: Africans (Gabon and Cameroon), Asians (Vietnam), and Europeans (Germany and UK). False discovery rate (FDR; Benjamini and Hochberg) correction was used to correct for multiple hypothesis testing. All p-values reported in the main text are uncorrected p-values. All FDR-corrected p-values are indicated as q-values or indicated accordingly. Spearman rho correlation was used for all correlations. One-tailed Binomial test was used to ask whether the slopes of genome size reduction by CI were consistent across bacterial genera.

Transfer events between Africa, Europe and Asia

To quantify the occurrence and directions of host-switch/transfer events, we used stochastic character mapping based on MCMC (*37*) implemented as SIMMAP in "phytools" R package (*33*). We applied the character mapping on (i) marker-based and MAG-based trees of adults in this study (five countries), (ii) marker-based trees of adults after adding public metagenomes (23 countries), (iii) marker-based trees of adults only using public metagenomes (19 countries), and (iv) marker-based and MAG-based trees of children in this study (three countries). Briefly, we treated "host regions" (i.e., "Africa", "America", "Asia", and "Europe") as bacterial traits, inferred ancestral character states on bacterial phylogeny using the equal rates model, repeated this 100 times, and calculated the average number of character changes and direction of host transfer events. Because larger trees will give a greater number of character changes on average, we divided the number of transfer events for each category by the total number of transfer events of that microbial tree. Occurrence of transfer events from Africa to Asia and from Africa to Europe were combined as "Africa to rest". The same aggregation was applied to the two other categories; "Asia to rest" and "Europe to rest". Correlations between occurrence of transfer events and CI were based on Spearman rho correlation. Pairwise comparisons between categories of transfer events were based on Wilcoxon rank sum test.

### References (supplemental information)

24. G. Zeller, J. Tap, A. Y. Voigt, S. Sunagawa, J. R. Kultima, P. I. Costea, A. Amiot, J. Böhm, F. Brunetti, N. Habermann, R. Hercog, M. Koch, A. Luciani, D. R. Mende, M. A. Schneider, P. Schrotz-King, C. Tournigand, J. Tran Van Nhieu, T. Yamada, J. Zimmermann, V. Benes, M. Kloor,

C. M. Ulrich, M. von Knebel Doeberitz, I. Sobhani, P. Bork, Potential of fecal microbiota for early-stage detection of colorectal cancer. *Mol. Syst. Biol.* **10**, 766 (2014).

25. P. Ferretti, E. Pasolli, A. Tett, F. Asnicar, V. Gorfer, S. Fedi, F. Armanini, D. T. Truong, S. Manara, M. Zolfo, F. Beghini, R. Bertorelli, V. De Sanctis, I. Bariletti, R. Canto, R. Clementi, M. Cologna, T. Crifò, G. Cusumano, S. Gottardi, C. Innamorati, C. Masè, D. Postai, D. Savoi, S. Duranti, G. A. Lugli, L. Mancabelli, F. Turroni, C. Ferrario, C. Milani, M. Mangifesta, R. Anzalone, A. Viappiani, M. Yassour, H. Vlamakis, R. Xavier, C. M. Collado, O. Koren, S. Tateo, M. Soffiati, A. Pedrotti, M. Ventura, C. Huttenhower, P. Bork, N. Segata, Mother-to-infant microbial transmission from different body sites shapes the developing infant gut microbiome. *Cell Host Microbe*. **24**, 133–145.e5 (2018).

**Supplemental figures**

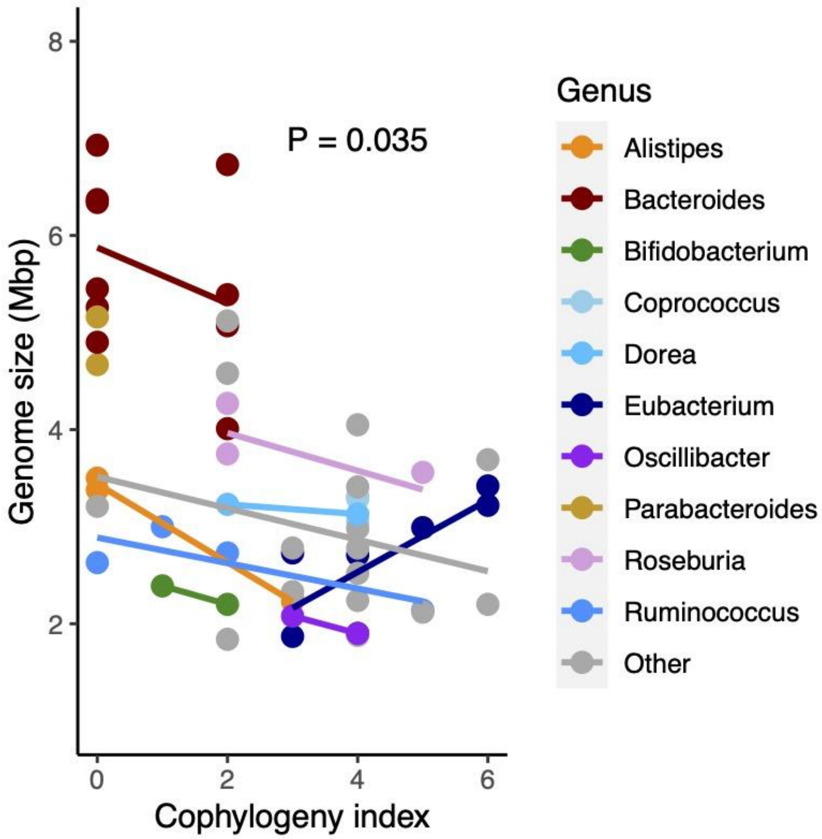


**Figure S1.** Correlation between cophylogeny index and genome size by microbial genus. Genera that contain two or more species in the list of the 56 adult taxa are highlighted in color. Genera that contain only one species are shown in gray. The p-value is based on a Binomial test: seven of eight genera show the expected relationship (decreasing genome size with great CI).

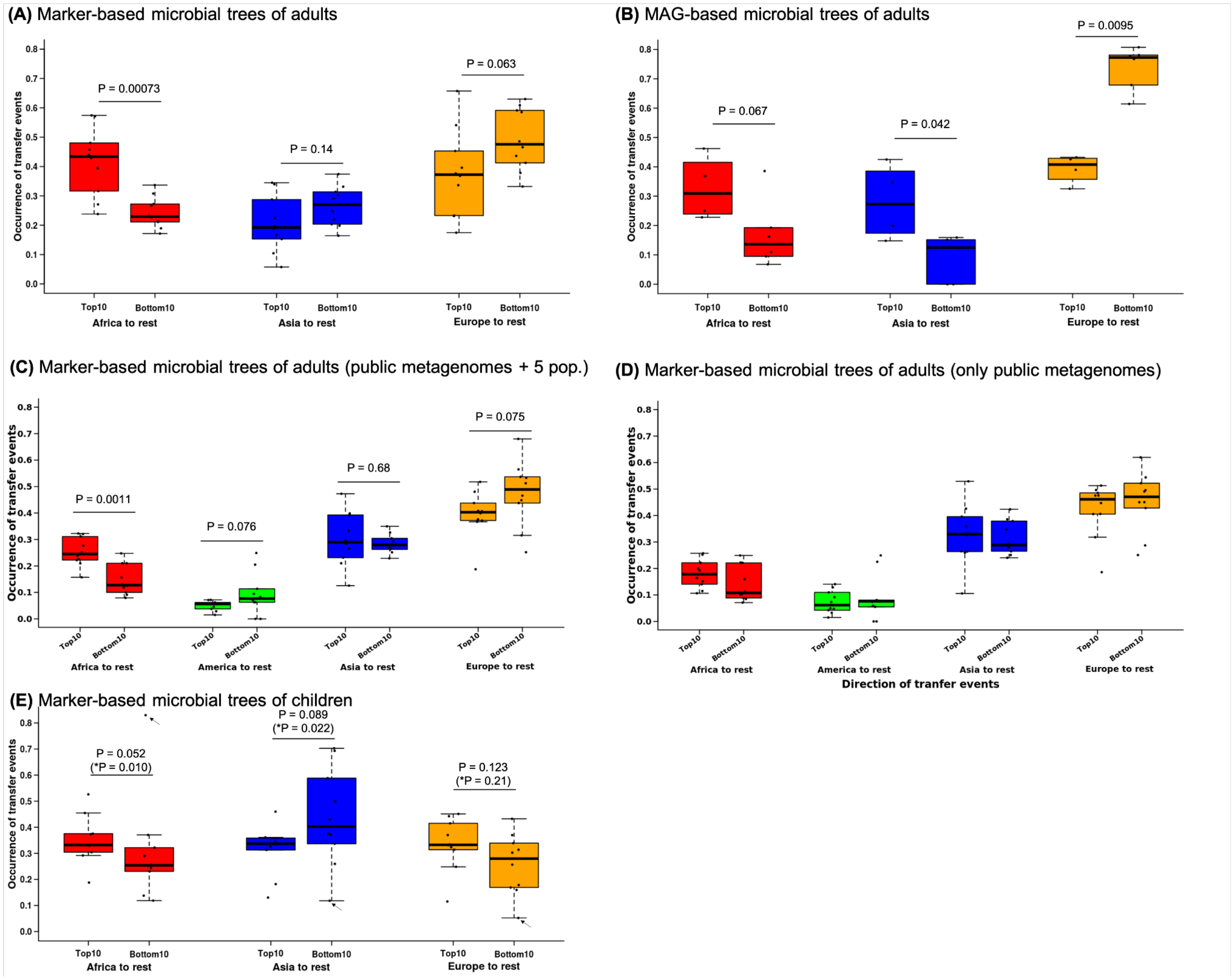


**Figure S2**. Boxplots comparing occurrence of transfer events between sampling regions for the top 10 and bottom 10 taxa in adults identified by the codiversification tests (Table S5). Results of stochastic character mapping on microbial trees based on (A) marker genes including adult metagenomes from five countries, (B) MAGs including adult metagenomes from five countries, (C) marker genes including adult human metagenomes from this study and the public dataset, (D) marker genes including adult human metagenomes from only the public dataset, and (E) marker genes including child metagenomes from three countries. The p-values are based on the Wilcoxon rank sum test. P-values with and without outlier (indicated by *, Grubbs test p = 0.0006) are shown for panel D.

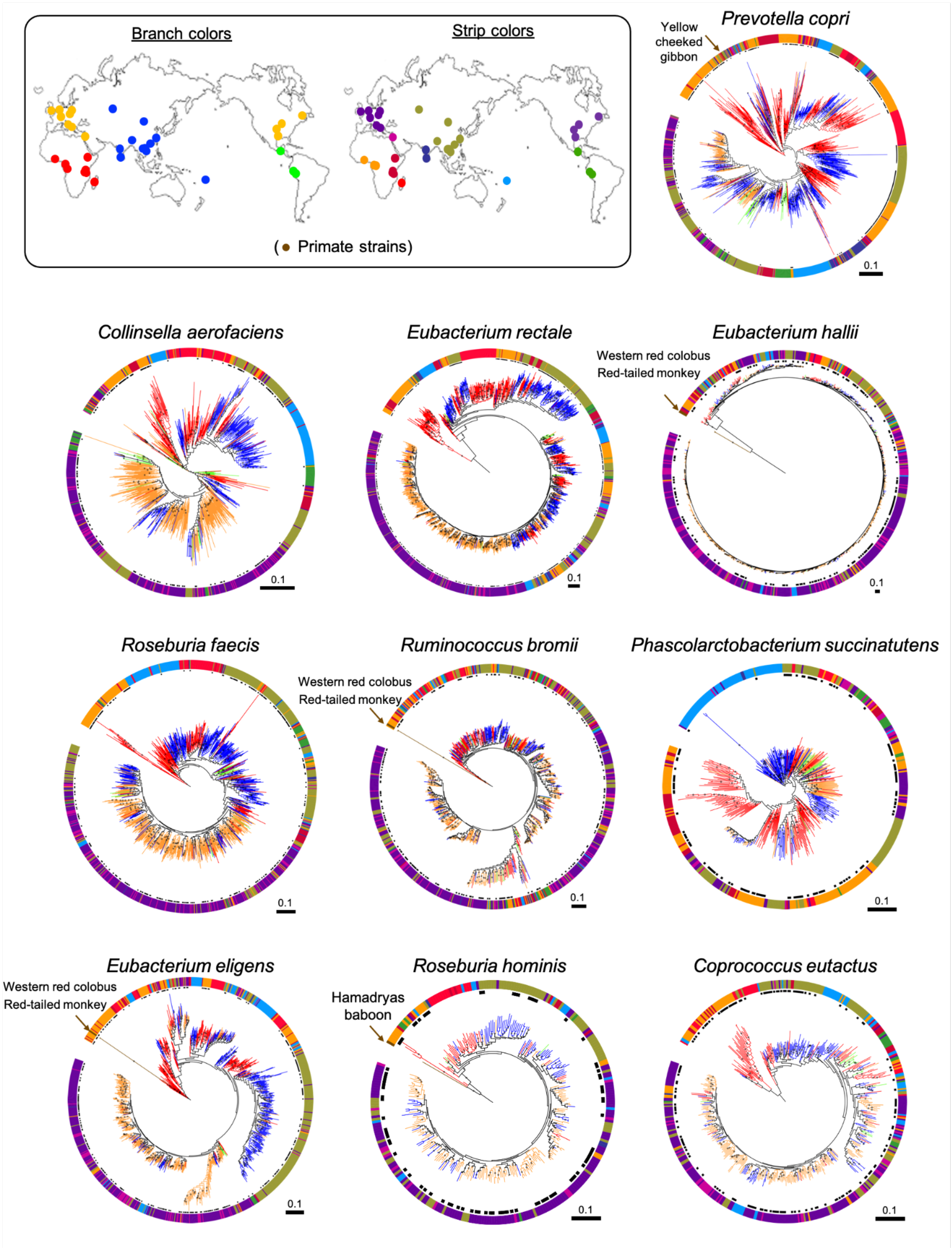


**Figure S3.** Microbial phylogenies of the top 10 taxa (highest CIs) with addition of public metagenomes. Colors of the branches and outer colorstrip indicate the host genetic structure estimated from sampling locations (*10*). Arrows indicate strains from Primate hosts. The black dots next to the outer colorstrip are samples from the initial analyses with five populations with paired fecal metagenomes and human genomes (*i.e.*, Gabon, Cameroon, Vietnam, Germany, and UK). Bootstrap values ≥50% are shown. All trees were rooted at the midpoint. The scales are substitutions per site.

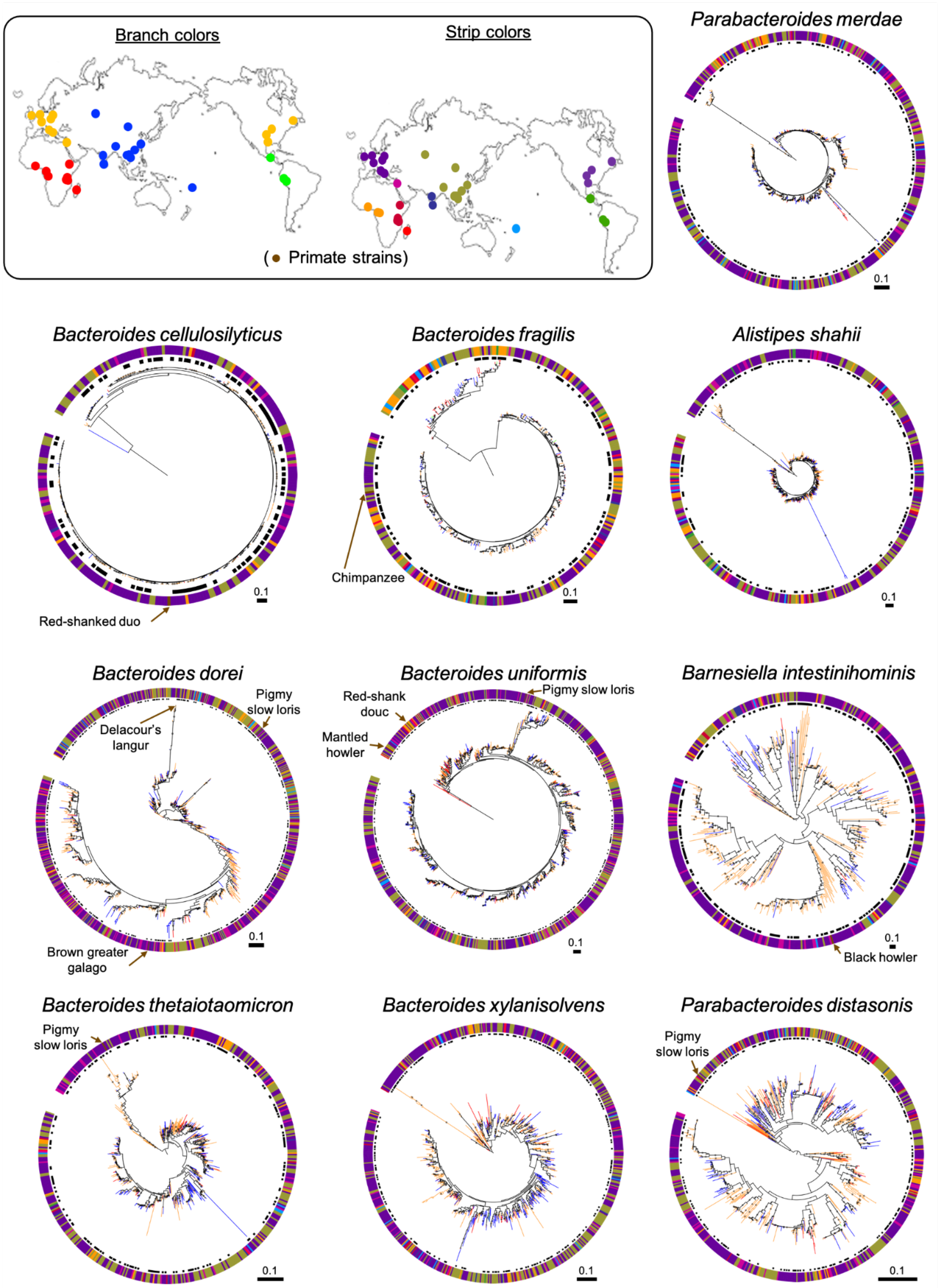


**Figure S4.** Microbial phylogenies of the bottom 10 taxa (lowest CIs) with additional public metagenomes. Colors of the branches and outer colorstrip indicate the host genetic structure estimated from sampling locations (*10*). Arrows indicate strains from Primate hosts. The black dots next to the outer colorstrip are samples from the initial analyses with five populations with paired fecal metagenomes and human genomes (*i.e.*, Gabon, Cameroon, Vietnam, Germany, and UK) where we tested for codiversification. Bootstrap values ≥50% are shown. All trees were rooted at the midpoint. The scales are substitutions per site.

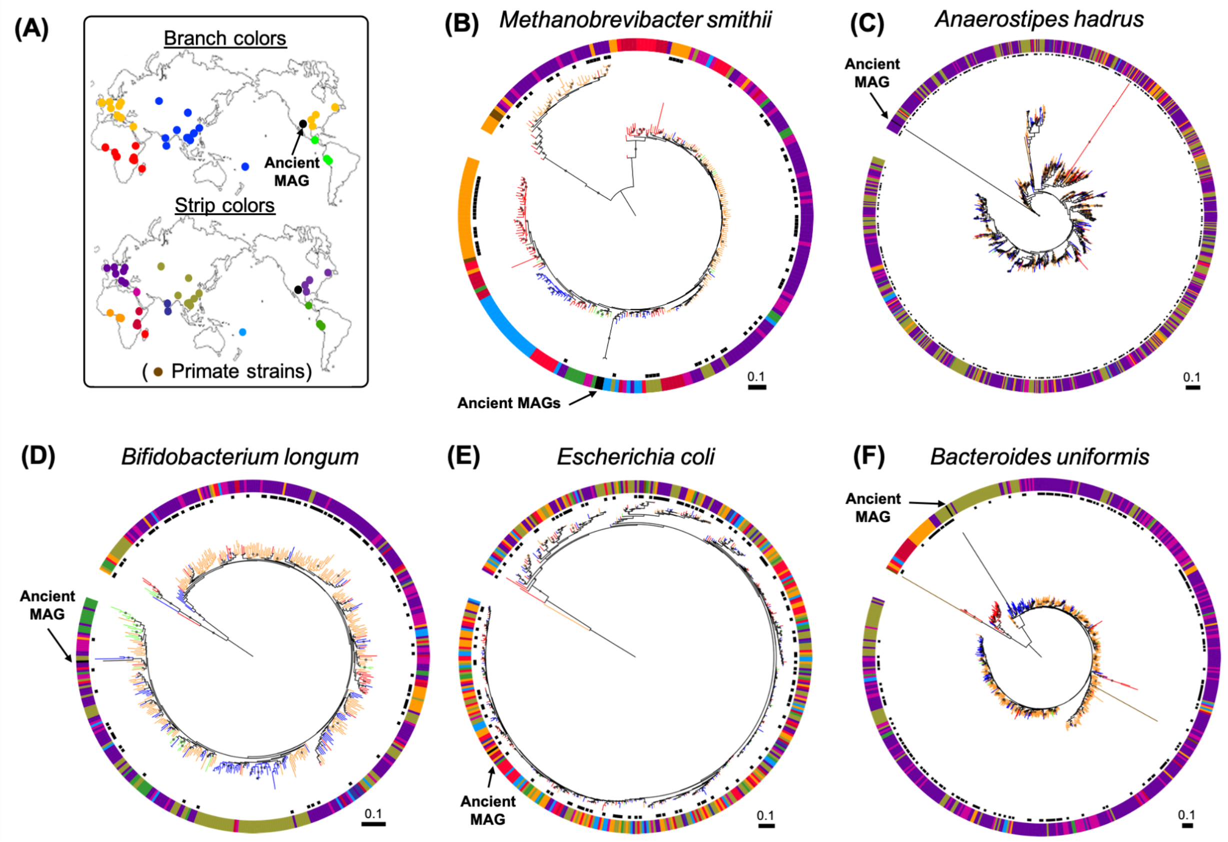


**Figure S5.** Microbial phylogenies that represent ancient MAGs from Native Americans. (A) The colors of the branches and outer colorstrip indicate the host genetic structure estimated from sampling locations (*10*). (B-F) Five microbial species that represent ancient MAGs. Arrows indicate strains from paleofeces of Native Americans. The black dots next to the outer colorstrip are samples from the initial analyses with five populations with paired fecal metagenomes and human genomes (*i.e.*, Gabon, Cameroon, Vietnam, Germany, and UK). Bootstrap values ≥50% are shown. All trees were rooted at the midpoint. The scales are substitutions per site.

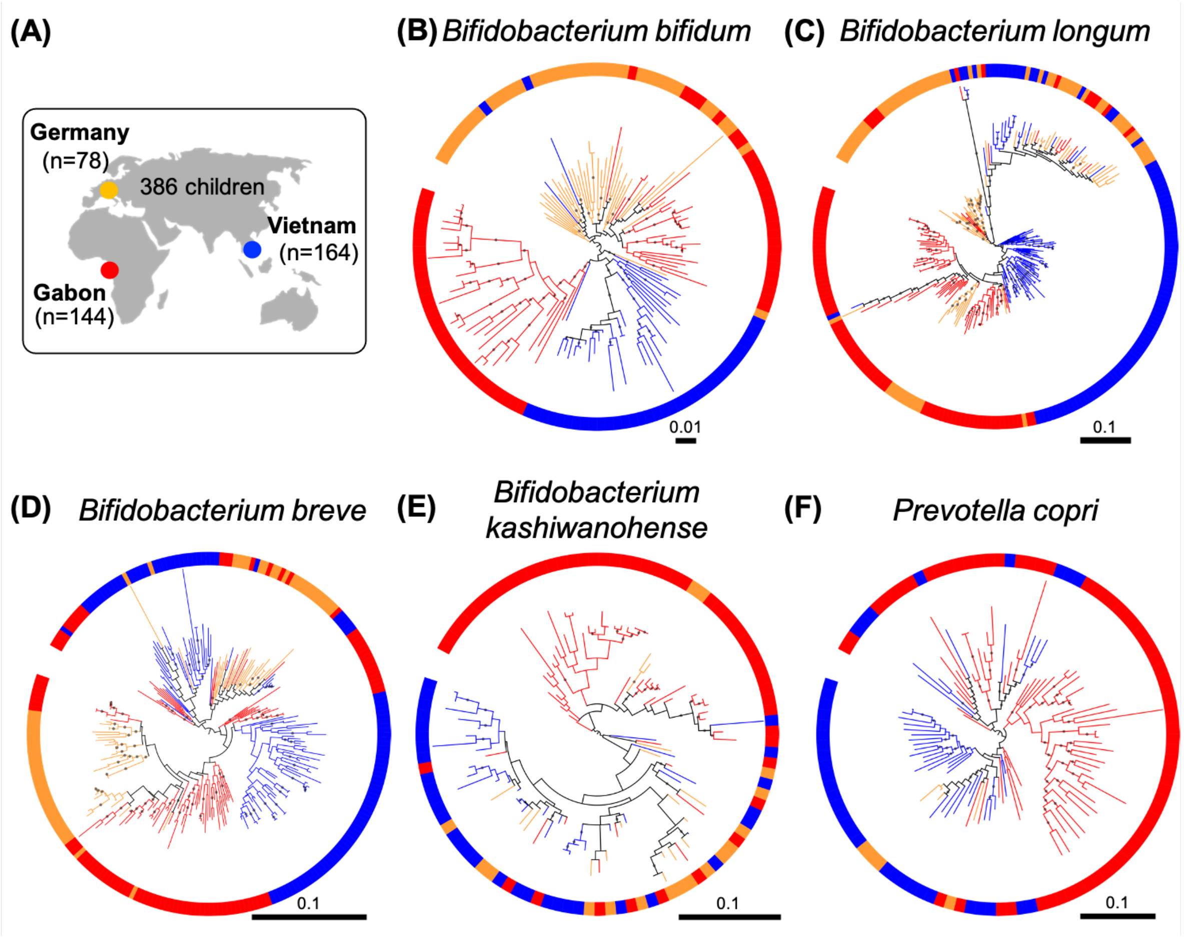


**Figure S6.** Examples of microbial phylogenies in child metagenomes. (A) A map of sampling locations and sample size are shown. Colors in the figure correspond to the map. (B-E) Microbial trees of four species of Bifidobacterium spp. that showed evidence of codiversification. (F) A microbial phylogeny of *P. copri* in children that also showed evidence of copylogeny in adults (Fig. 1D). Bootstrap values ≥50% are shown and the phylogenies are rooted at the midpoint. Codiversification test results for all 20 taxa are in Table S6. The scales are substitutions per site.

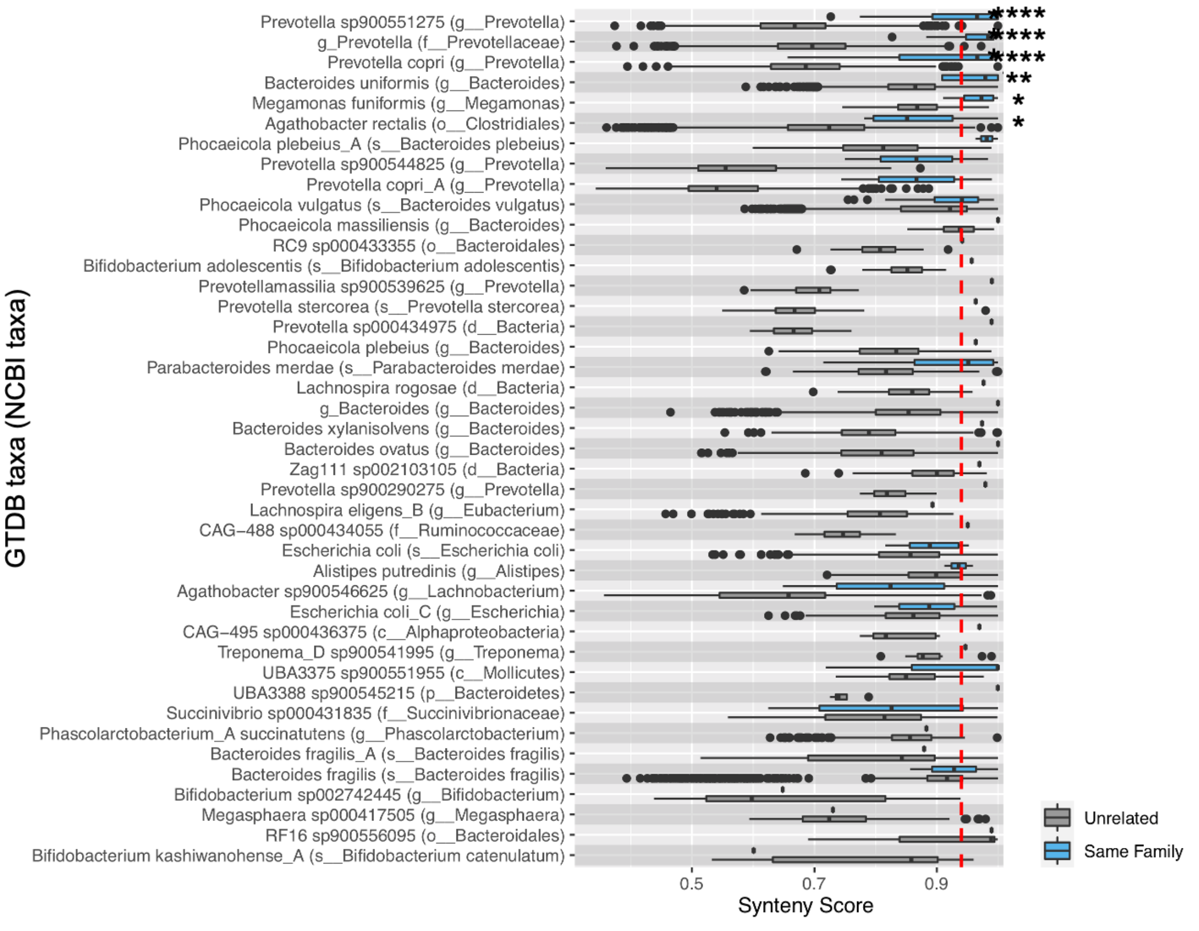


**Figure S7.** Strain sharing assessed within related (blue) and unrelated (gray) mother-child pairs using SynTracker. The relatedness of strains per species within related and unrelated mother-child pairs is shown as Synteny Score. Species listed are those identified in both related and unrelated mother-child pairs. Dashed red lines indicate the pairwise comparison thresholds for strain sharing events (Synteny Score = 0.96). Stars correspond to q-values (corrected Wilcoxon-Mann-Whitney test; * = $q < 0.05$; ** = $q < 5x10^{-3}$; **** = $q < 5x10^{-7}$).

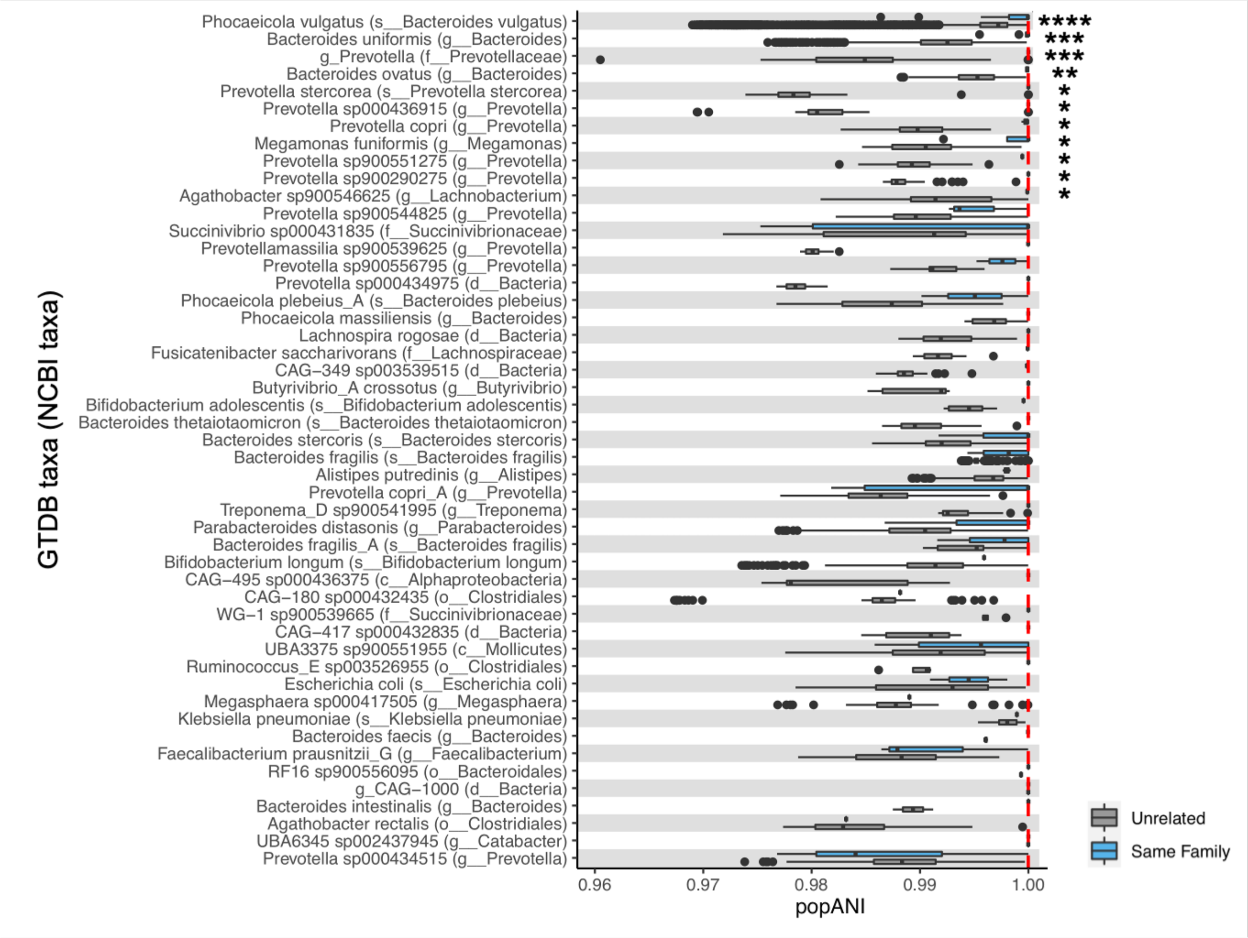


**Figure S8.** Strain sharing assessed within related (blue) and unrelated (gray) mother-child pairs using inStrain. The relatedness of strains per species within related and unrelated mother-child pairs is shown as popANI. Species listed are those identified in both related and unrelated mother-child pairs. Dashed red lines indicate the pairwise comparison thresholds for strain sharing events (pop ANI = 0.99999). Stars correspond to q-values (corrected Wilcoxon-Mann-Whitney test; * $q < 0.05$, ** = $q < 5x10^{-3}$; *** = $q < 5x10^{-4}$; **** = $q < 5x10^{-10}$).

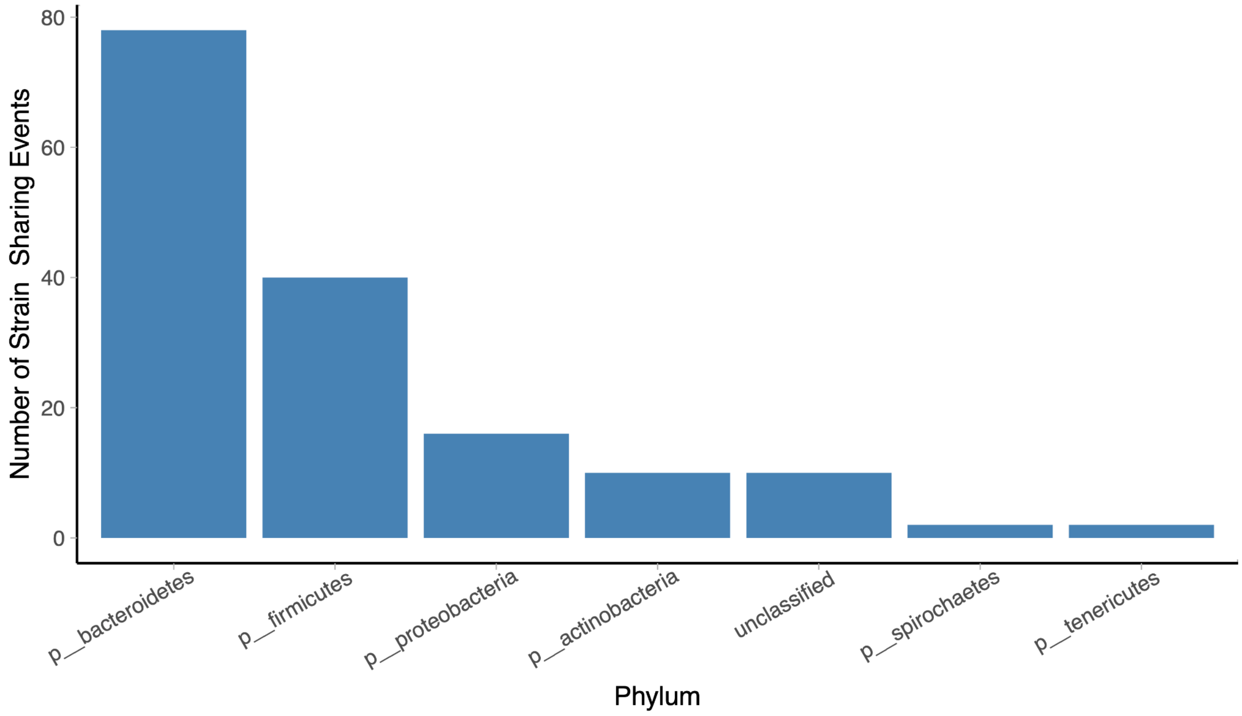


**Figure S9.** Phyla of strains shared between related mother-child pairs, unrelated mother-child pairs sampled in the same location, and unrelated mother-child pairs sampled in different locations.
